# Detecting convergent adaptive amino acid evolution

**DOI:** 10.1101/513010

**Authors:** Carine Rey, Vincent Lanore, Philippe Veber, Laurent Guéguen, Nicolas Lartillot, Marie Sémon, Bastien Boussau

**Affiliations:** Univ Lyon, Université Claude Bernard Lyon 1, ENS de Lyon, CNRS UMR 5239, INSERM U1210, LBMC, F-69007, Lyon, France.; Univ Lyon, Université Claude Bernard Lyon 1, CNRS UMR 5558, LBBE, F-69100, Villeurbanne, France.

**Author notes:** Those authors contributed equally (Carine Rey, Vincent Lanore). Those authors contributed equally (Marie Sémon, Bastien Boussau).

**Keywords:** convergent evolution, genomics, molecular evolution, coding sequences, phylogenetics, probabilistic models

## Abstract

In evolutionary genomics, researchers have taken an interest in identifying in the genomes substitutions that subtend convergent phenotypic adaptations. This is a difficult question to address, because genomes contain billions of sites, many of which have substituted in the lineages where the adaptations took place, and yet are not linked to them. Those extra substitutions may be linked to other adaptations, may be neutral, or may be linked to mutational biases. Furthermore, one can think of various ways of defining substitutions of interest, and various methods that match those definitions have been used, resulting in different sets of candidate substitutions. In this manuscript we first clarify how adaptation to convergent phenotypic evolution can manifest itself in coding sequences. Second, we review methods that have been proposed to detect convergent adaptive evolution in coding sequences and expose the assumptions that underlie them. Finally, we examine their power on simulations of convergent changes, including in the presence of a confounding factor.

## INTRODUCTION

It is difficult to replicate experiments when we study evolutionary biology. However, one can benefit from natural replicates that have arisen through time and across taxa, because different lineages have been subjected independently to the same “experimental” conditions. In such cases, lineages have adapted independently to the same environmental constraints. In evolutionary genomics in particular, researchers have taken an interest in identifying in the genomes substitutions that subtend those adaptations. This is a difficult question to address, because genomes contain billions of sites, many of which have substituted in the lineages where the adaptations took place, and yet are not linked to them. Those extra substitutions may be linked to other adaptations, may be neutral, or may be linked to mutational biases. Furthermore, one can think of various ways of defining substitutions of interest, and various methods that match those definitions have been used, resulting in different sets of candidate substitutions (1–3). The purpose of our manuscript is first to clarify the definition of convergent adaptive amino acid evolution by examining the processes that can create it. Second, we review the existing methods to detect convergent amino acid evolution and expose the assumptions that underlie them. Third we examine their power on simulations of convergent changes, including in the presence of a confounding factor.

### Defining convergent adaptive amino acid evolution

It is useful to first think about adaptive evolution before tackling convergent adaptive evolution. Adaptive genomic evolution is expected to occur when constraints on the phenotype change, which alters the selective pressures at some sites in the genome. Individuals with mutations that provide an increased fitness in the new environment will have a reproductive advantage, so these mutations will increase in frequency and could eventually fix. The fixation of one or some of these mutations in turn can change the selective pressures operating on the sites of the genome: because of epistatic interactions, mutations that were e.g. advantageous can now become even more advantageous, neutral, or deleterious (4). The characteristics of this fitness landscape have an impact on how likely convergent adaptive evolution will be. First, if a particular site always provides the highest fitness increase when the phenotype changes, convergent evolution at this very site is more likely. But if different sites provide similar fitness advantages, different substitutions may fix in different lineages, making convergent evolution less likely. Subsequently if the fixation of one or the other of these early mutations changes the fitness landscape, convergent evolution of further late-fixing mutations is less likely. These intuitive considerations should make it clear that a good mechanistic model of convergent evolution needs to consider the entire genome at a time, along with the fitness landscape, to take into account all the dependencies between sites. For computational reasons, and because fitness landscapes are only rarely studied experimentally (4), such a model is currently out of reach. Instead, each site is typically modelled independently of the others, and simplifying assumptions are made: for instance, fitness landscapes only depend on the phenotype, and not on the lineage under consideration. From now on, we will consider such simplified models of convergent genomic evolution.

In this article we propose to define convergent evolution through the comparison of coding sequences across species. Coding sequences offer a window into where the mutation process and the selective process meet, since some non-synonymous mutations that change amino acids will be strongly counter selected while other synonymous mutations, which keep coding for the same amino acid, will be neutral or weakly counter-selected. The simplest codon models consider one site at a time and allow two classes of substitutions: the non-synonymous and the synonymous ones, and assume that synonymous substitutions provide a proxy for the rate of fixation of neutral substitutions, while all non-synonymous substitutions have the same rate of fixation, which depends on selection efficacy (5,6). More sophisticated codon models distinguish between different amino acid changing substitutions, and assume that biochemical similarity between amino acids affects how interchangeable they are in a protein. Such models use amino acid fitness profiles—which we simply call *amino acid profiles* in the rest of the manuscript (Figure 1) (7). Some of the richest models allow different parameters for different sites of a protein (8,9). Overall, codon models provide a sufficient framework to define convergent adaptive amino acid evolution.

**Figure 1:**
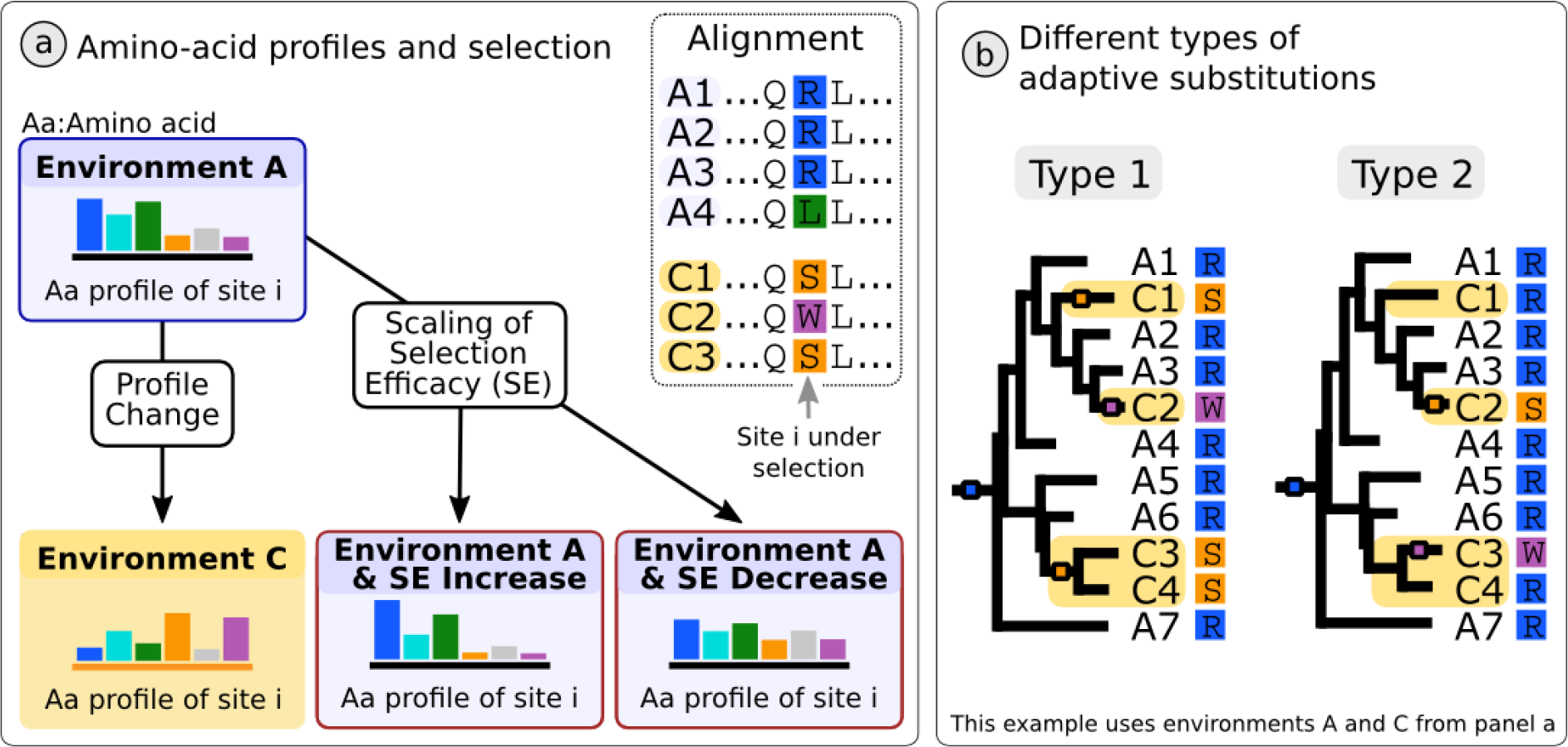
Categories of adaptive and non-adaptive convergent amino acid evolution.Left: At a particular position in a protein, some amino acids provide better fitness than others. This is represented by colored bars for 6 amino acids, the bigger the bar the higher the fitness. In the ancestral environment A, amino acids blue and green provide the highest fitness, whereas in the convergent environment C those are amino acids orange and purple. Increasing the selection efficacy makes the profiles more pointed, while decreasing it makes them more flat, but the amino acid relative rank does not change. Right: Species with the convergent phenotype are named C* and species with the ancestral phenotype are named A*. Substitutions are small boxes on the branches. We distinguish two types of adaptive convergent substitutions: Type 1 are substitutions that occur systematically on the branch where the phenotype changes, at the transition between Ancestral and Convergent environments (A to C). Type 2 are substitutions that occur on later branches (e.g. in the branch leading to C3).

In a simple model that considers one codon at a time, adaptive evolution can manifest itself by increasing the selective pressure, changing its nature, but not by decreasing it. Indeed, relaxations of the pressure can hardly be described as adaptive; for instance although it might be slightly advantageous for subterranean animals to stop producing opsin proteins in an environment without light (10) the relaxation of selection on the sequence of opsins and other visual genes is more likely to be associated to a lack of selective pressure on the sequences of those proteins, which have become useless, rather than positive selection to optimize the allocation of resources and stop producing a small amount of useless proteins (11). Increases of the pressure would mean that, in the amino acid profile at a given codon, the same amino acids that provided high fitness before the change still provide high fitness after the change, but even more so, while amino acids providing low fitness before the change now provide even worse fitness (Figure 1, left panel, “Scaling of Selection Efficacy”). It has become more important for the organism to have particular amino acids at this position. This could be associated to a lifestyle where the function of the protein has become more important than it was. In addition, we will study changes in the nature of the selective pressure that manifest themselves by a change between two amino acid profiles, which will be referred as ancestral and convergent in the following (Figure. 1, left panel “Profile Change”). In this case we expect that different amino acids will provide high fitness to the organism before and after a phenotype change. In a given condition, it may be that a single amino acid provides much more fitness than all other amino acids, or that several amino acids provide an equivalent fitness.

Furthermore, we can distinguish two cases, depending on whether the substitution happens on the same branch as the phenotypic change, or on later branches (Figure 1, right panel). In the former case, type 1, it may be that the substitution caused or was highly related to adaptation to the convergent phenotype; perhaps it was even necessary for the organism to have the convergent phenotype. In the second case, type 2, the substitution may still provide a fitness advantage in the new phenotype, but it is not necessary; perhaps it provides a fitness advantage in the convergent phenotype, given that pre-existing important substitutions have already fixed.

### Detecting adaptive convergent amino acid evolution

Several methods have been designed to detect convergent adaptive amino acid evolution. We list them below and attempt to predict their relative strengths and weaknesses, in particular their capacity to predict type 1 and type 2 convergent adaptive substitutions. All of the following methods have been designed to detect some type of profile change, so we expect that they will do much better to detect convergent profile changes than to detect convergent increases or decreases in selection efficacy.

#### Method based on topological incongruencies

The “topological” method is an early attempt to look for an indirect effect of convergent sequence evolution, based on an observation first made on the prestin gene (12) and later systematized in a genome-scale study (1–3). When a particular site has evolved convergently in several lineages, it will display the same or similar amino acids in those lineages, and not in lineages with a different phenotype. As a result, for this site a phylogeny in which lineages with the convergent state are grouped together will be more likely than the true species phylogeny. This approach involves constructing the species topology and a “convergent” topology where species with the convergent phenotype are grouped together. Then, each site can be tested for which topology it prefers, the true species phylogeny or the convergent phylogeny, by comparing the likelihoods of the two trees for this site. This method is capturing a byproduct of convergent evolution, and not its mechanism, hence it is difficult to know precisely what type of substitution this method can work with. Presumably both type 1 and type 2 substitutions can be detected.

#### Methods looking for independent substitutions to the same amino-acid

The most intuitive method, the “identical” method, looks for independent substitutions to the exact same amino acid in all clades with the convergent phenotype (13,14). It therefore assumes that a particular amino acid has a much better fitness than all other amino acids at this particular position of a protein. In practice, it relies on ancestral sequence reconstruction to infer the amino acids present before each convergent transition and make sure that the transition of interest occurred on the branch where the phenotypic transition occurred. By design, it is very conservative because it can only detect sites where a single particular amino acid is much more fit than others, which fixed with a type 1 substitution (Figure 2).

**Figure 2:**
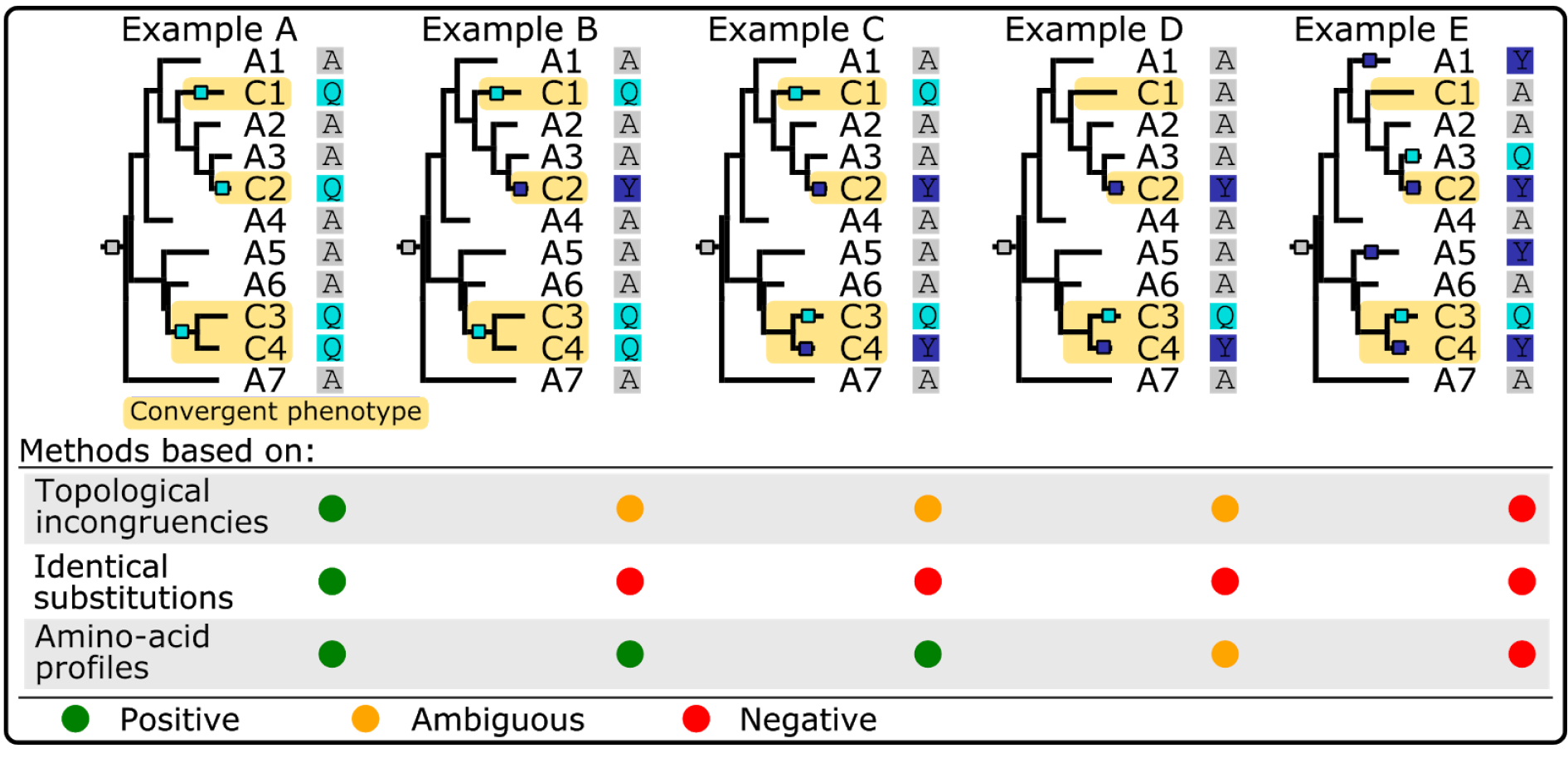
Cartoon examples of sites included or not in the target sites of each convergence detection tool. The tree topologies and species are the same in all examples. Species with the convergent phenotype are named C*, those with the ancestral phenotype A*; the transitions between ancestral and convergent phenotype occur where the subtrees become shaded in yellow. Colored squares on the branches of the phylogeny indicate substitution events, with the color corresponding to the arriving state. In Example A, every time the phenotype changes, a substitution occurs to amino acid Q (type 1 substitutions to a single amino acid). This is an ideal case for the methods based on identical substitutions, and should be detectable by all methods. Example B shows a profile change, whereby 2 different amino acids, Q and Y, have good fitness in the convergent case. All methods but the Identical may detect such changes, although this depends on how different the ancestral and the convergent profiles are (17). Example C is similar to Example B except that some substitutions occurred after the phenotype has changed (type 2 substitutions), not simultaneously with the phenotype change. Example D is similar to Example C except that the amino acid change only occurred 3 times out of 4: this makes it more controversial, and harder to detect. But if the change in profile is strong enough, profile methods should be able to detect it. Example E shows a case where the evolution of the site does not seem to correlate with the convergent/ancestral state of the species. We don’t expect the methods to detect such a site, but some such sites will nevertheless come out as false positives.

An extension of this method, the “expectation” method of Chabrol et al. (15), also called msd, looks for sites with a high convergence index. This convergence index is the expected number of substitutions to a particular amino acid in lineages with the convergent phenotype. Interestingly, and contrary to the other methods presented here, this method does not assume that the lineages where the phenotypic changes occurred must be known. Instead, it is enough to have phenotypic annotations for extant species only. It is unclear whether it will be very conservative or not: on one hand it will detect only sites where a particular amino acid is found in most species with the convergent phenotype, as in the “identical” method, but on the other hand this convergence could apply to a subset of the species with the convergent phenotype only, an advantage compared to methods based on amino-acid profiles (see below). Both type 1 and type 2 substitutions can be detected by this method, but type 2 substitutions will get a higher convergence index than type 1 substitutions and may therefore be better detected.

#### Methods based on amino-acid profiles

“Profile methods” are methods aiming to detect pressure changes, whereby different amino acids will provide good fitness in the ancestral and convergent phenotypes. The simplest of them is the “Multinomial” approach, which compares the amino acid frequencies in extant species with the ancestral phenotype with the amino acid frequencies in extant species with the convergent phenotype using a simple Chi2 test for multinomial distributions (16). This approach has not been used in the literature to our knowledge and suffers from a major drawback in that it fails to account for the phylogenetic structure of the data. However, we chose to include it in our tests as it provides a baseline against which the other more sophisticated methods can be tested. Both type 1 and type 2 substitutions can be detected by this method.

Other profile methods include PCOC, diffsel and TDG09, which belong to a family that we loosely call “mechanistic methods”, because they combine a phylogenetic approach with amino acid fitness profiles.

The “PCOC” method (17) models convergent evolution at the amino acid level, without taking into account the codon level. It combines the “profile” idea by attributing different equilibrium frequencies, which act as fitnesses, to the 20 amino acids, before and after the phenotypic changes, with the One Change (OC) model. OC assumes that sites involved in the convergent adaptation must have undergone a substitution on the branches where the adaptation took place. Detection of sites having undergone convergent evolution is obtained by comparing the likelihoods of two models, one where convergent evolution is assumed complete, with the change in equilibrium frequencies and enforced substitutions on all branches where the phenotype changed, and another where evolution is homogeneous across all branches. Amino acid profiles are not estimated, but are drawn from pre-existing distributions that have been estimated on large collections of alignments (18). Both type 1 and type 2 substitutions can be detected by PCOC, but with different power: the OC component of PCOC expects only type 1 substitutions, but the PC component can accommodate both type 1 and type 2 substitutions.

The TDG09 model (19) is similar to PCOC in that it handles amino acid sequences, but it focuses on profile changes and does not include the OC component. Further, it estimates the profiles based on each site of the alignment. To do so, it builds two profiles, one for the species with the ancestral phenotype, and one for the species with the convergent phenotype. Amino acids with a count at or under 2 are considered absent, and all absent amino acids are assigned a 0.0 frequency in the profile vector. To detect sites undergoing convergent adaptive evolution, a likelihood ratio test is performed between a model that assumes a single profile across the entire tree, or two profiles for the ancestral and convergent parts of the tree. Both type 1 and type 2 substitutions can be detected by such methods.

Finally, diffsel (20) is similar in spirit to TDG09 but works at the codon level. In this codon model, mutations occur at the DNA level, and selection occurs at the amino acid level. Selection is modelled as a fitness profile of 20 amino acid fitnesses. Convergent sites are characterized by a systematic change from an ancestral amino acid fitness profile to a different amino acid profile on all branches where the phenotype changed. Both type 1 and type 2 substitutions can be detected by such methods. In this manuscript, their detection has been based on summarizing Bayesian MCMC output (see methods).

All those methods look for particular patterns that can be detected from the comparison of sequences in a range of species and that are suggestive of adaptive convergent amino acid evolution. However, these patterns can also be generated by neutral processes or by mutational biases.

### Non-adaptive convergent amino acid evolution

Even in the absence of selection, some amount of convergent amino acid evolution is expected, if only because there are only 20 possible amino acids. Further, the structure of the genetic code and the characteristics of the mutation process (e.g. that transitions are more frequent than transversions) all contribute to making some amino acid substitutions more likely than others, and therefore increase the probability that they will be convergent.

In addition, fixation biases could create patterns resembling convergent adaptive evolution. In particular, GC-biased gene conversion (bGC) is a fixation bias that favours G or C alleles over A or T alleles and is widespread across the tree of life (21,22). It is most intense in regions of the genome that recombine frequently, and will have a stronger effect over time in species with large effective population sizes and short generation times. Those two characteristics have appeared independently several times in the tree of life. Because of bGC, one can expect to detect convergent changes to GC alleles in the species sharing these characteristics, even without any adaptive value to having GC alleles instead of AT at those positions. This phenomenon seems to be strong enough to affect single gene phylogenies in birds (23,24), and may be an important driver of non-adaptive convergent sequence evolution.

Convergent global relaxations of selection could also create patterns that look like adaptive convergence. If the phenotypic change is linked to a genome-wide decrease in selection efficacy, e.g. through a decrease in the effective population size (25), mutations that used to be counter selected become tolerated. Combined with the structure of the genetic code, the same substitution could occur in lineages undergoing the decrease in selection efficacy.

Finally epistatic interactions between sites in the genome or within a protein can create non-adaptive convergent amino acid evolution (26). The same mutation at a particular site can occur in independent lineages simply because by chance sites that are in epistatic interactions with it happen to be in the same state in those lineages. The mutation therefore fixes not because of an adaptation to a new environment, but because of the states of interacting sites. Such non adaptive convergence is more likely in closely related lineages than in distant lineages.

## RESULTS AND DISCUSSION

Some of the methods presented above have been implemented in several software packages (Table 1). In this manuscript, we test these software packages on simulated data along with methods we have re-implemented ourselves. We evaluate the power of the methods in 3 cases, (i) a convergent profile change, (ii) a convergent increase or decrease in selection efficacy (iii) and a combination of the above two, whereby a convergent profile change occurs simultaneously with a scaling of selection efficacy. To achieve this scaling, we set a selection efficacy parameter which is the product of 2 parameters, the population size (Ne) and the selective pressure (S) (it is also called a scaled selection coefficient). In the following, we will refer to this value by NeS, a composite parameter whose increase (resp. decrease) can be interpreted as e.g. a genome-wide increase of population size, or a site-wise increase of selective pressure, or both. We choose to use NeS=4 as the reference value, which produces alignments similar to empirical alignments according to a range of statistics (Figures S3-S6).

**Table 1:**
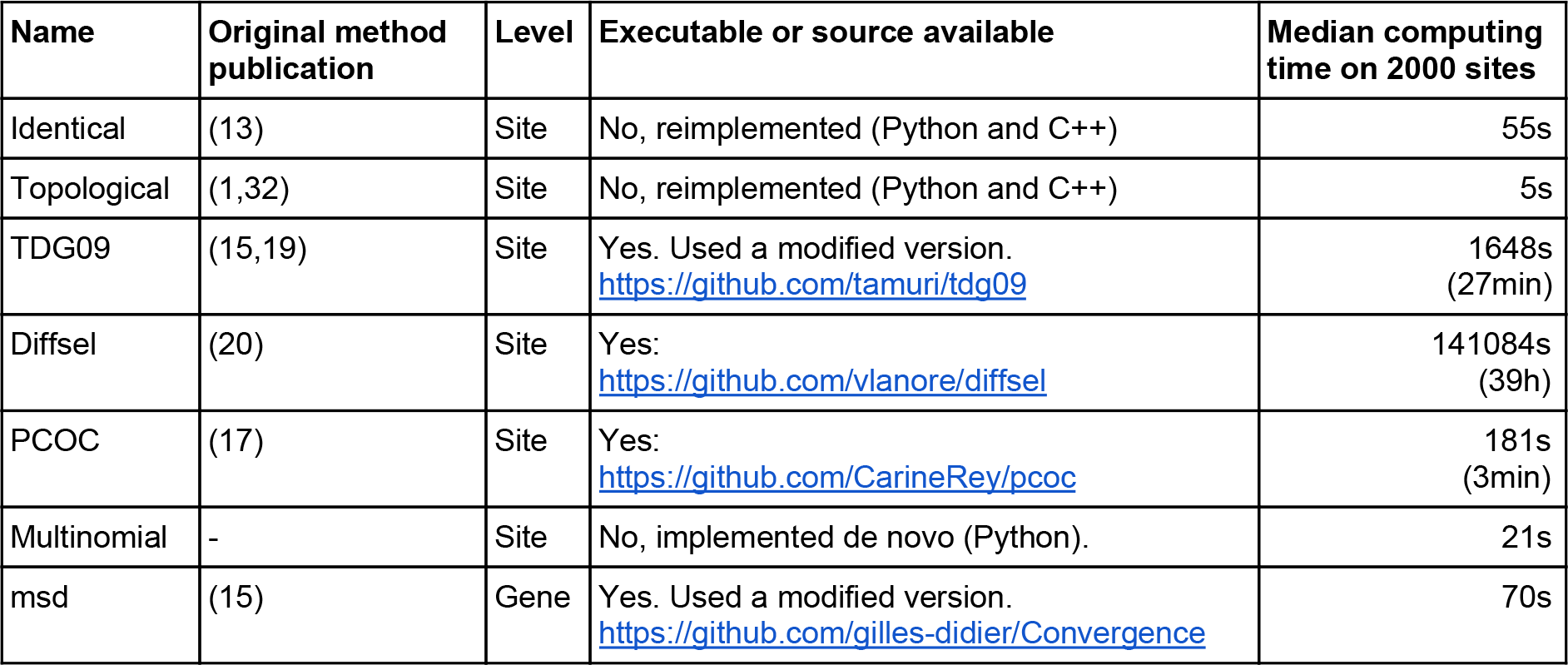
Summary of the methods used in the pipeline

| Name | Original method publication | Level | Executable or source available | Median computing time on 2000 sites |
| --- | --- | --- | --- | --- |
| Identical | (13) | Site | No, reimplemented (Python and C++) | 55s |
| Topological | (1,32) | Site | No, reimplemented (Python and C++) | 5s |
| TDG09 | (15,19) | Site | Yes. Used a modified version.<br>https://github.com/tamuri/tdg09 | 1648s<br>(27min) |
| DiffSel | (20) | Site | Yes:<br>https://github.com/vlanore/diffsel | 141084s<br>(39h) |
| PCOC | (17) | Site | Yes:<br>https://github.com/CarineRey/pcoc | 181s<br>(3min) |
| Multinomial | - | Site | No, implemented de novo (Python). | 21s |
| msd | (15) | Gene | Yes. Used a modified version.<br>https://github.com/gilles-didier/Convergence | 70s |

We tested the methods on 4 empirical phylogenies with different size, depth and number of transitions (13,27,28), (Figure S1).

In case 1, “Convergent profile change”, selection efficacy remains constant but the amino acid profile changes between the ancestral and convergent conditions. To simulate this case (Figure 3, Figure 4 top row), we change the amino acid profile in the convergent clades and we keep the same global NeS along the tree. The results are presented in Figure 3 for NeS=4, and for the 4 empirical phylogenies.

**Figure 3:**
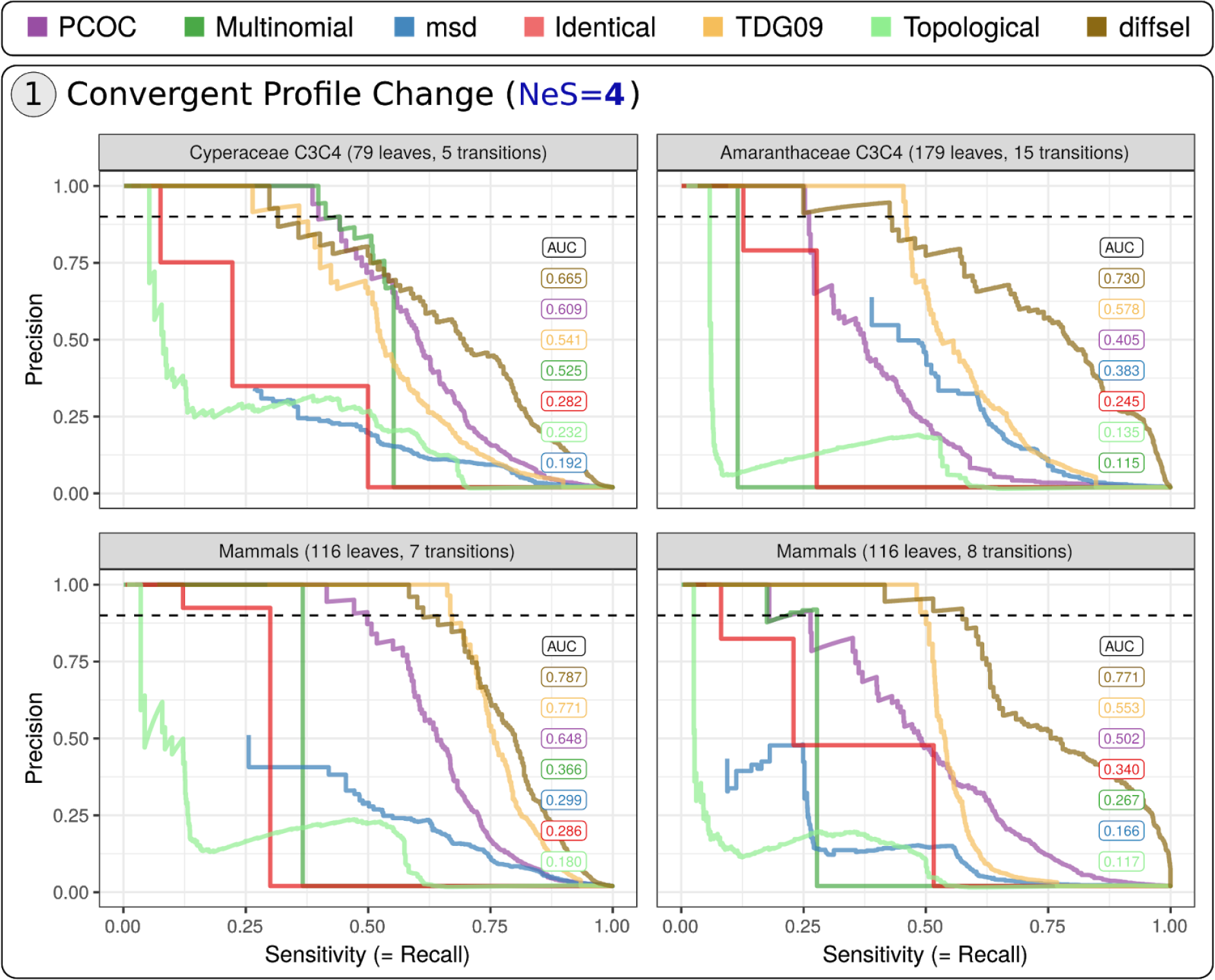
Detection of sites undergoing convergent profile change by different methods. Simulations are performed with constant selection efficacy (NeS=4). Each panel corresponds to one empirical phylogeny, with convergent transitions placed as in Figure S1. The trade-off between sensitivity and precision is presented for each method, assuming that 2% of the sites are convergent in the sequences (color code indicated on the top of the figure). The dashed line highlights 90% precision. Area Under the Curves (AUC) ranked from best to worst are presented in the top right corner for each panel, with the same color code as the precision-recall curves.

**Figure 4:**
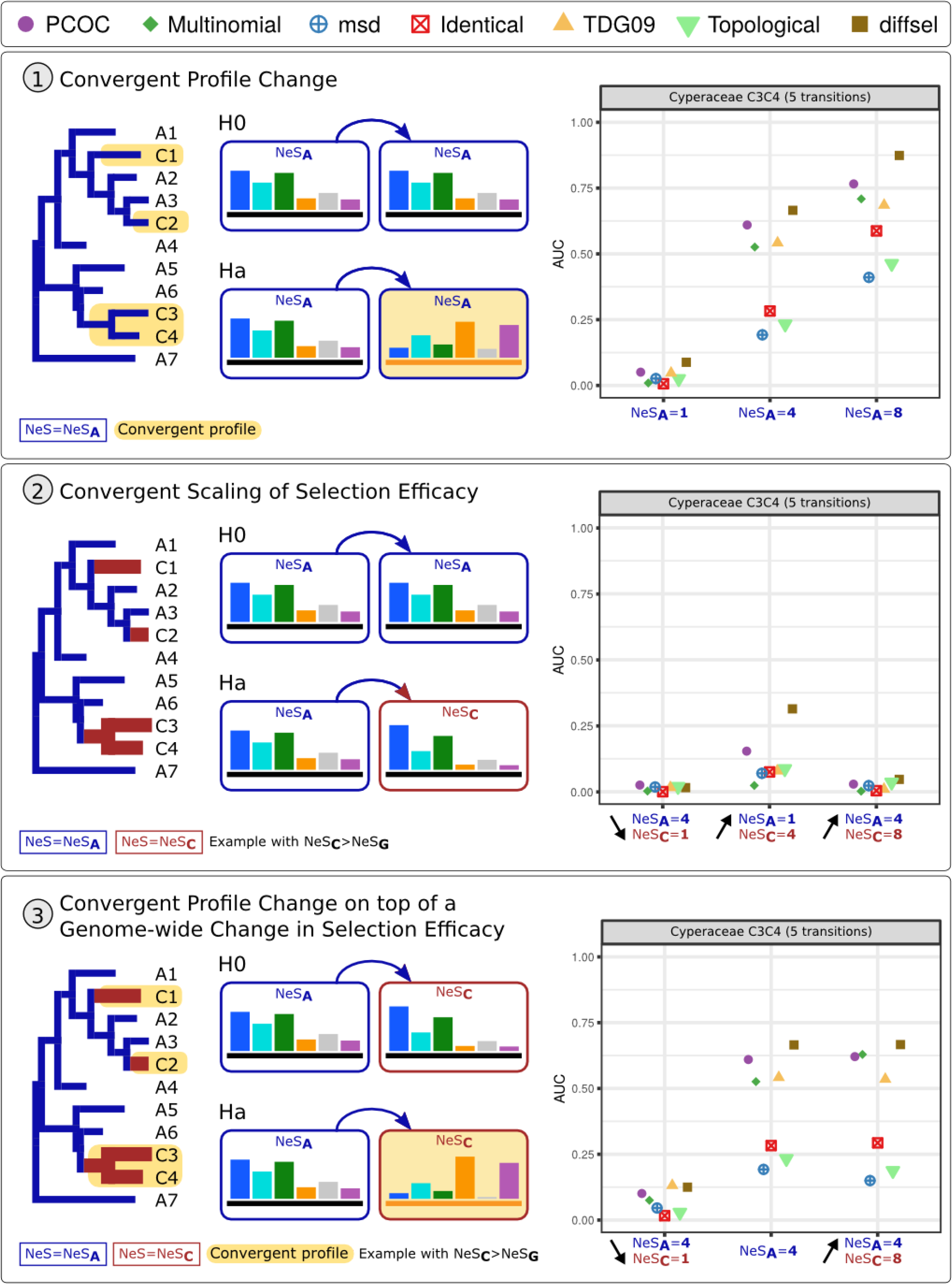
Overview of the simulations and AUC values for the Cyperaceae tree. The trees, convergent clades and symbols are as in Figure 1. Three kinds of adaptive convergent cases have been simulated: 1) a convergent profile change, 2) a convergent scaling of selection efficacy, and 3) a convergent profile change combined with a selection efficacy scaling. The genome-wide selection efficacy (NeS_A_) remains the same in 1) and is changed to a convergent selection efficacy (NeS_c_) in Ha 2) and Ha 3). The black arrows (2 and 3) indicate if selection efficacy increases or decreases in convergent clades. AUC values are calculated based on precision-recall curves such as presented in Figure 3 (top row: AUC values for NeS_A_=4 in the case 1 correspond to the top left of Figure 3).

Profile methods perform better than the other methods in the 4 phylogenies, and among them, diffsel dominates the benchmark according to AUC values (Figure 3). The sensitivity at 90% precision is not as easily interpretable as AUC because the curves are very rugged; TDG09, PCOC and diffsel seem to dominate this metric, with a different order depending on the tree. Surprisingly, the simplistic Multinomial method performs well on the Cyperaceae tree, competing with the TDG09 and PCOC in terms of its sensitivity at 90% precision. The relative ranks of PCOC, Multinomial and TDG09 vary depending on the tree, which may be attributable to differences in the number of convergent transitions and in the relative size of the convergent clades. For instance, we suspect that PCOC’s performance is degraded when the number of convergent transitions increases, because by design it looks for sites with convergent changes in all the convergent clades, not just a subset of them. TDG09 shows the opposite trend, with better performance when the number of transitions increases. The Topological, Identical and msd approaches typically perform worse, but the AUC rank of msd is volatile. The low sensitivity of Identical and msd is expected as those methods can only detect convergent substitutions to a particular amino acid, not to an amino acid profile. Overall, these results are qualitatively congruent with previously published simulations obtained with simpler settings and fewer methods (17). However, the precisions and sensitivities observed here are much worse than those reported in (17), because simulations do not use the Profile Change with One Change model, which enforces substitutions on transition branches.

Note that diffsel, which performs well in our experiments, is also the most expensive method computationally by several orders of magnitude (Table 1). Other methods may be preferable for large datasets unless extensive computing resources are available. The better performance of profile methods may be due to their fitting the simulation conditions better. However, it is unclear how we could have simulated convergent evolution realistically without using mutation-selection models that use profiles of amino acid frequencies. In the end, this indicates that profile methods may perform better on empirical data as well; apart from diffsel which always comes out first, the variability of the AUC ranks among trees however indicates that using several methods on a data set is recommended.

We then studied the performance of the methods for a wider range of genome-wide selection efficacies, focusing on the Cyperaceae tree (see Figure S7 for the three other trees). The top row of the Figure 4 represents AUC values for the Cyperaceae tree, for NeS=1, 4 and 8, corresponding to values for weak, medium and high selection efficacy respectively, all of which produce alignments with realistic properties (Figures S3-S6). As expected, the methods are most accurate when NeS is high (NeS = 8), and the performance collapses when selection is not efficient (NeS = 1). In other words, it should be extremely difficult to detect convergent molecular evolution in species with small Ne, or for sites under weak selective pressure.

In case 2, “Convergent scaling of selection efficacy” (Figure 4, middle row), the same amino acid fitness profile is used along the whole tree for a given site, but NeS is changed in convergent clades (from NeS_A_ to NeS_C_) in Ha simulations. It is important to note that a NeS variation implies the modification of the fitness of each amino acid in the profile but not of its rank (see Figure 1, left). We made 3 runs, two with an increase and another with a decrease of NeS in convergent clades. Overall, methods perform poorly at detecting selection efficacy scaling, with the exception of the NeS_A_=1 to NeS_C_=4 case where PCOC and diffsel detect a small number of sites.

By the 2 previous cases, we saw that methods can detect adaptive convergent sites under two conditions: they have undergone a profile change and they are under moderate to high selective pressure. But the methods cannot detect profile changes when selection efficacy is low, and also fail to detect scalings in selection efficacy alone.

Finally, case 3 introduces a confounding factor. Here we assume a genome-wide scaling of selection efficacy on top of which convergent sites undergo profile changes (Figure 4, bottom row), and we try to detect those latter sites. This is modeled by a selection efficacy scaling from NeS**_G_** to NeS**_C_**in both convergent (Ha) and non-convergent (H0) sites, plus an amino acid profile change in Ha. We tried both to decrease (left panel) or increase (right panel) the selection efficacy in the convergent clades and compared the results to the situation obtained when selection efficacy is constant. With a decreased selection efficacy in convergent clades, the methods’ performances deteriorate compared to the reference simulation. With an increased selection efficacy in convergent clades, the performances remain roughly the same. In other words, a decrease in selection efficacy (for instance due to a decrease in Ne) coinciding with convergent transitions has a negative impact on the detection of convergent profile changes, but an increase has very little impact.

Those results reveal the performance of existing methods to detect two different types of convergent amino acid evolution on simulated data, in isolation or combined with each other. The simulations have been performed with complex models of sequence evolution, parameterized so as to generate data sets that resemble empirical data on a few test statistics. However, some key assumptions underlying those models are clearly unrealistic: first, each site is simulated independently of the other ones. It would be useful to incorporate epistatic constraints in our simulations as those will increase the number of non-adaptive convergences (26). Such a model has been proposed (26,29), but can only be used on proteins whose structure has been solved, and requires assuming that the structure is constant across all the tree.

Second, although it is an important part of the model, the phenotype is here considered in an extremely naive fashion. In particular, we have made no effort to incorporate a distribution of fitness effects, whereby different sites would contribute differently to the phenotype under consideration, and therefore to the fitness (30). Using such a distribution would be key to understanding why some sites, those of large effect, undergo convergent evolution while others, with smaller effects, do not. It could also indicate to users what effect sizes are large enough to be detected in a given experimental setting, and what effect sizes are just too small to be detected.

Third, several known confounding factors have not been simulated. In particular, we have not incorporated biased Gene Conversion in our simulations, and we have not incorporated population-level processes that would allow polymorphisms to cross speciation events (incomplete lineage sorting, ILS) and would increase the levels of polymorphisms present at the tips of the trees.

With these caveats in mind, our simulations show that existing methods are much better at detecting convergent profile changes rather than convergent selection efficacy scalings. Further, detection of convergent profile changes is improved when selection efficacy is high, possibly because this increases the frequency of type 1 substitutions. They also show that model-based methods, that explicitly rely on profiles, perform better than other methods.

Moving forward, we can think of three complementary directions for improving methods aiming to detect convergent adaptive evolution in amino-acid sequences. In all cases, they will be based on profile methods anchored in a mechanistic modeling of sequence evolution. As a first direction, we need to complement models of sequence evolution so that, in addition to profile changes we can also detect accurately changes in selection efficacy, and distinguish those adaptive processes from confounding factors such as biased gene conversion and ILS. Further anchoring the model in population genetics theory may allow interpreting detected sites in terms of the fitness advantage they provide. As a second direction, we need to improve the computational efficiency of model-based inference. This should be a major concern here, because data sets are getting larger every year; algorithmic or mathematical developments will probably be necessary to fit such complex models onto large data sets. In this respect, one intriguing result of this study is the performance of the Multinomial method. This simplistic method ignores nearly everything of the complexities of codon models of sequence evolution, and yet achieves a performance that rivals them in some conditions. Correcting the Multinomial method for phylogenetic inertia could provide even better performances, and it may be possible to make it able to further improve it while keeping excellent speed. Finally, we have only tested the methods’ ability to detect individual convergent sites; some methods (e.g. msd) can also employ a statistical procedure to detect convergent genes by combining site-wise evidence. Alternatively, TDG09 has a procedure to control its false positive rate, and diffsel estimates parameters based on entire alignments, not single sites. All those features have not been tested but are crucial for application to real data, in particular for application to genome-wide data sets. Future analyses will have to investigate these aspects.

## METHODS

### Simulation of alignments of coding sequences

We simulated coding sequences using bppseqgen (31) under (heterogeneous) mutation-selection models, which belong to the “mechanistic” family of methods tested in this work. Mutation-selection models are codon models that combine mutations at the DNA level with amino acid fitness vectors, so that selection operates only at the amino acid level. Our mutation-selection models were complemented by a parameter indicating the efficacy of selection, NeS. In our mutation-selection model, NeS controls the flatness of the amino acid profiles (see supplementary section 1). With a high NeS, the profiles are very peaked, and with a low NeS, very flat. We investigated the impact of different NeS values, in homogeneous models, where the same NeS is applied to all the branches (Figure 4, first row), and in heterogeneous models, where different NeS are used for the branches in the ancestral and convergent parts of the tree.

We performed several types of simulations. Simulation settings are described in the results section (Figure 4). For each simulated codon position, one or two profiles are selected randomly in our set of 263 non-redundant profiles and one or two NeS values are chosen. One profile and one NeS value is used for the ancestral branches, and the others for convergent subtrees.

### Methods to detect adaptive convergent evolution

In order to compare results across methods, it was necessary to standardize their output. See the supplementary material for details.

### Pipeline and implementation of the methods

The results in this paper were obtained using an all-in-one pipeline that encompasses simulations, detection and post-simulation analysis, including the generation of the plots used for Figure 3 and Figure 4. The pipeline itself was implemented in OCaml using bistro (https://github.com/pveber/bistro), a library to build statically-typed reproducible workflows. Special attention was paid to reproducibility, in particular by following the guidelines given in (33). Instructions to reproduce our results are given in the supplementary material.

The implementations of the methods used in the pipeline are as follows:

- The Multinomial method has been implemented de novo in Python as well as the Identical and Topological methods which additionally use executables from the bppsuite (31). They are available via the pipeline.
- The TDG09 implementation we used is a slightly modified version of the one available on github (see Table 1) where multithreading has been removed to avoid multithreading-related problems. Results should be identical to github version. In addition, a script available in the pipeline repository was written to adapt input alignments and trees to TDG09 expected formats.
- For diffsel, we used an optimized version of the original implementation that is faster but implements the same model. The implementation we used is available on github (see Table 1). In addition, we use a different approach to establish MCMC convergence. The original method compares two MCMC chains using the tracecomp program from the PhyloBayes suite (34). Instead, we run only one chain, use the Raftery and Lewis’s Diagnostic implemented in the R package coda (v0.19-1) (35) after 200 iterations to estimate the number of necessary iterations, then run as many iterations as 120% of the estimated number, and finally perform the same diagnostic to check convergence.
- We used the github version of PCOC (see Table 1) as is.
- Regarding msd, we used a version modified by the author so as to output a p-value for all sites, which we needed to compute scores.

The experiments performed for this paper—*i.e.* the whole pipeline with 2000 sites for each hypothesis times 12 hypotheses times 4 trees—took five days to run on a 24-core virtual machine. Computation times observed during this run for individual detection methods are given in Table 1. Note that most of the computing time for the whole pipeline is spent in diffsel tasks, which are a lot more costly to compute than other methods.

## CONCLUSION

We have reviewed existing definitions of convergent amino acid evolution. We have built upon them to distinguish between convergent amino acid profile changes and convergent scalings in the efficacy of selection. When existing methods are tested on simulated data in a range of conditions, probabilistic methods that rely on models of sequence evolution detect convergent profile changes better than other methods. However, none of them performs well at detecting convergent scalings of selection efficacy, and they perform poorly when selection efficacy is low over the entire tree. Improved models would allow distinguishing between different types of convergent evolution, and should use mathematical and algorithmic tricks to improve computational efficiency.

## Data, code and materials

Our pipeline’s code is available at https://gitlab.in2p3.fr/pveber/reviewphiltrans. It contains everything required to reproduce our results. Detailed reproduction instructions are given in supplementary section 8. All intermediate data used to produce our results (~20Go) is temporarily available at ftp://pbil.univ-lyon1.fr/pub/lanore/ during the review process (and would be hosted on Dryad after acceptance).

## Authors’ contributions

CR, VL and PV built the pipeline for testing the methods. CR, VL, LG, NL and BB implemented some of the methods. NL estimated empirical profiles. CR, MS and BB performed statistical analyses. CR, VL, MS and BB wrote the manuscript. All authors read, commented on and approved the manuscript.

## Acknowledgements

We would like to thank G. Didier for providing a modified version of msd that can work on individual sites, and T. Latrille for providing the profiles and fruitful discussions. This work was performed using the computing facilities of the CC LBBE/PRABI. We would like to thank the French Institute of Bioinformatics - IFB CNRS UMS 3601 - (funded as part of Investissement d’avenir program managed by Agence Nationale pour la Recherche, contract ANR-11-INBS-0013) for providing life science data and tools, storage and computing resources on the IFB national service infrastructure in bioinformatics.

