## Supplementary material for "Detecting convergent adaptive amino acid evolution"

[*]: Those authors contributed equally

[§]: Those authors contributed equally

1/ Model of codon sequence evolution used in the simulations 2

2/ Tree topologies 3

3/ Selection of amino acid profiles used in the simulations 5

Step 1: Estimation of site-wise profiles based on mammalian alignments 5

Step 2: Selection of a subset of non-redundant Bloom profiles 6

4/ Choice of NeS used in the simulations 8

5/ Standardizing the output of detection methods 11

6/ Comparaison of the AUC values for the 4 phylogenies 13

7/ Impact of using separate alignments for diffsel 15

8/ How to reproduce our results 16

**References 17**

### 1/ Model of codon sequence evolution used in the simulations

We performed simulations using bppseqgen (1,2) under a codon model in which all sites are considered independent of each other, without rate heterogeneity across sites. This codon model accounts for site-wise amino acid preferences and for variations in the efficacy of selection. For a given site, it has two levels, with level 1 modelling DNA mutations between codons, and level 2 modelling the fixation of these mutations, similar to (3–5).

At level 1, mutations between codons are based on a Jukes-Cantor model (6), allowing only one nucleotide substitution per codon per instantaneous time.

At level 2, fixation probabilities are dependent on two parameters: the vector of amino acid frequencies, and the efficacy of selection NeS. If the frequencies of two amino acids A and B are $f_A$ and $f_B$, the fixation probability of the mutation from A to B is proportional to $-NeS\,\frac{log(f_A/f_B)}{1-(f_A/f_B)^{NeS}}$ as per Equation 2 in (7).

Finally, to fit with the usual modelling in phylogenetics, the resulting generator matrix is normalized so that there is one substitution per codon per unit of time at equilibrium.

#### 2/ Tree topologies

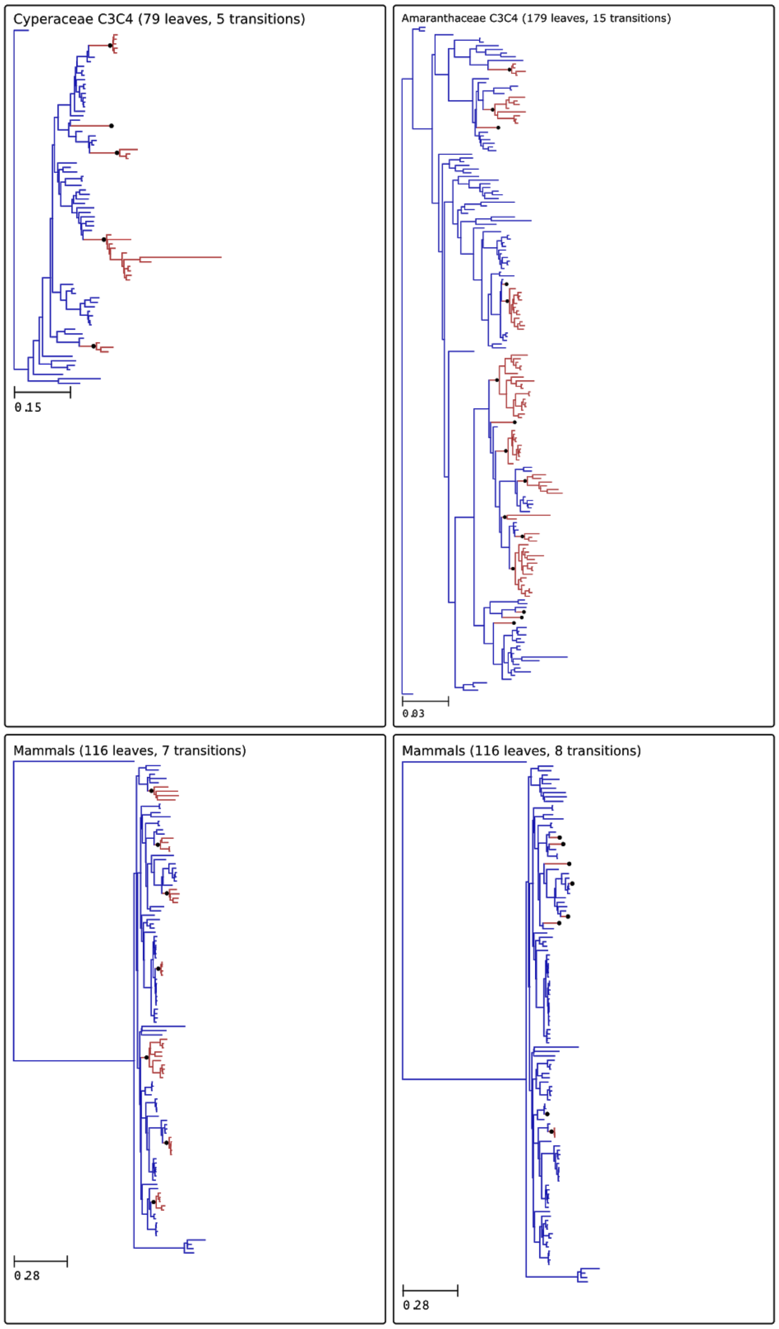


**<u>Figure S1:</u>** Overview of the 4 phylogenies. Convergent clades are in red. Black dots indicate branch with a convergent transition. Convergent transitions of the Cyperaceae (8) and the Amaranthaceae (9) phylogenies correspond to C3C4 transitions. For the mammals phylogenies (10), convergent transitions are placed on branches leading to xeric species for the 8 transitions tree. For the 7 transitions tree, convergent transitions have been placed unrelated to a specific phenotype, but homogeneously in the tree such as convergent clades are small subtrees. Tree topologies and convergent transition annotations are available at https://gitlab.in2p3.fr/pveber/reviewphiltrans.

### 3/ Selection of amino acid profiles used in the simulations

Amino acid profiles are vectors of frequencies for each of the 20 amino acids. These profiles are related to vectors of amino acid fitnesses (4). Several sets of amino acid profiles have been proposed in previous works. Our aim was to obtain a set of non-redundant and realistic amino acid profiles that would readily describe the frequencies of amino acids for a site of a protein alignment that has been subjected to a constant and homogeneous set of selective forces throughout its evolution. This criterion discards sets of profiles estimated from amino acid alignments (11), as those must contain sites that have undergone changing selective pressures during their history. We therefore relied on profiles that have been obtained during wet lab experiments involving mutagenesis and fitness measurements on viruses (12). We collected 1389 profiles by combining the profiles obtained in (12). However these profiles (named below “Bloom profiles”) are highly redundant (Figure S2-A), and therefore some filtering is necessary before using them to simulate changes in fitness profiles. We reasoned that a good subset of these profiles would be one that can fit most sites of empirical alignments well. To obtain this subset, we used a two step approach. First, we learned thousands of site-wise amino acid profiles from alignments of mammalian coding DNA sequences. Second, we estimated the set of Bloom profiles that best fits 95% of these sites. This 95% criterion is arbitrary but corresponds to our hypothesis that only a minority of sites have undergone a change in selective pressures during their evolution, and therefore could not be well fitted by a single Bloom profile.

#### Step 1: Estimation of site-wise profiles based on mammalian alignments

Starting from Orthomam v6 (13), 100 CDS were randomly sampled, for 37 placental mammals. The alignments were trimmed for misaligned regions and short frameshifts, yielding a total of 64721 aligned coding sites. To achieve faster computation, these were further subdivided into 5 sets of 20 genes. For each set, the alignments were concatenated, resulting in 5 independent multiple sequence alignment of approximately 13000 coding positions each (gene lists and alignments available…). For each alignment, the mutation-selection model of Rodrigue et al (5), such as implemented in phylobayes mpi version 1.8 (14) was run in 2 replicates, for approximately 4500 cycles. Counting a burnin of 500, these runs were used to compute the site-specific marginal posterior mean fitness profiles (64721 sites, 2 replicates).

#### Step 2: Selection of a subset of non-redundant Bloom profiles

For each of the 64721 profiles, we identified the amino acid profile from Bloom’s data that maximizes the multinomial likelihood. Finally, we chose the smallest set of Bloom profiles that provides ML fits for 95% of the 64721 profiles. This set contains 263 Bloom profiles. Amino acid frequencies of each profile are available in the gitlab repository of the pipeline and presented in Figure S2-B.

A

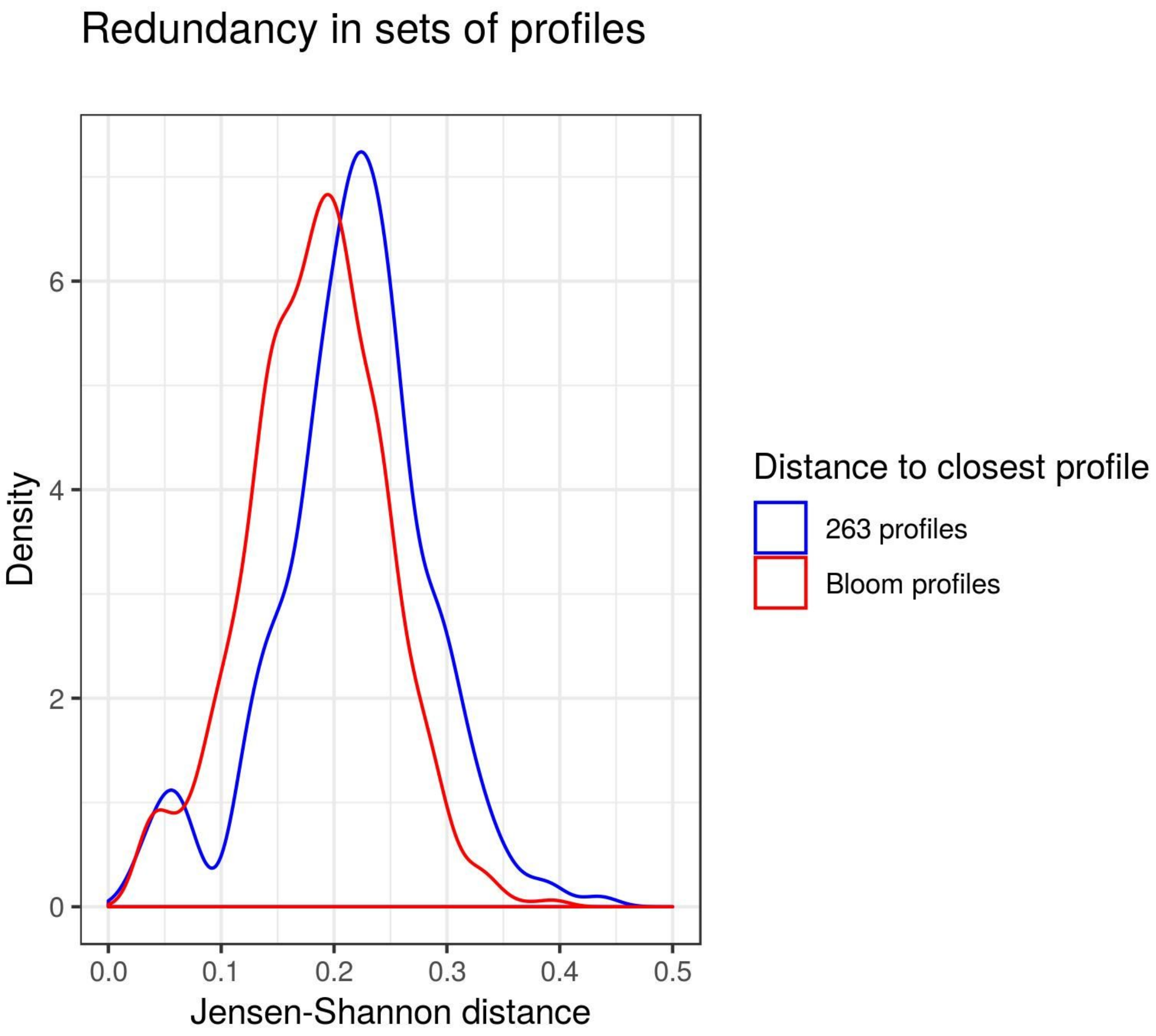
Redundancy in sets of profiles
Density
6
4
2
0
0.0
0.1
0.2
0.3
0.4
0.5
Jensen-Shannon distance
Distance to closest profile
263 profiles
Bloom profiles

B

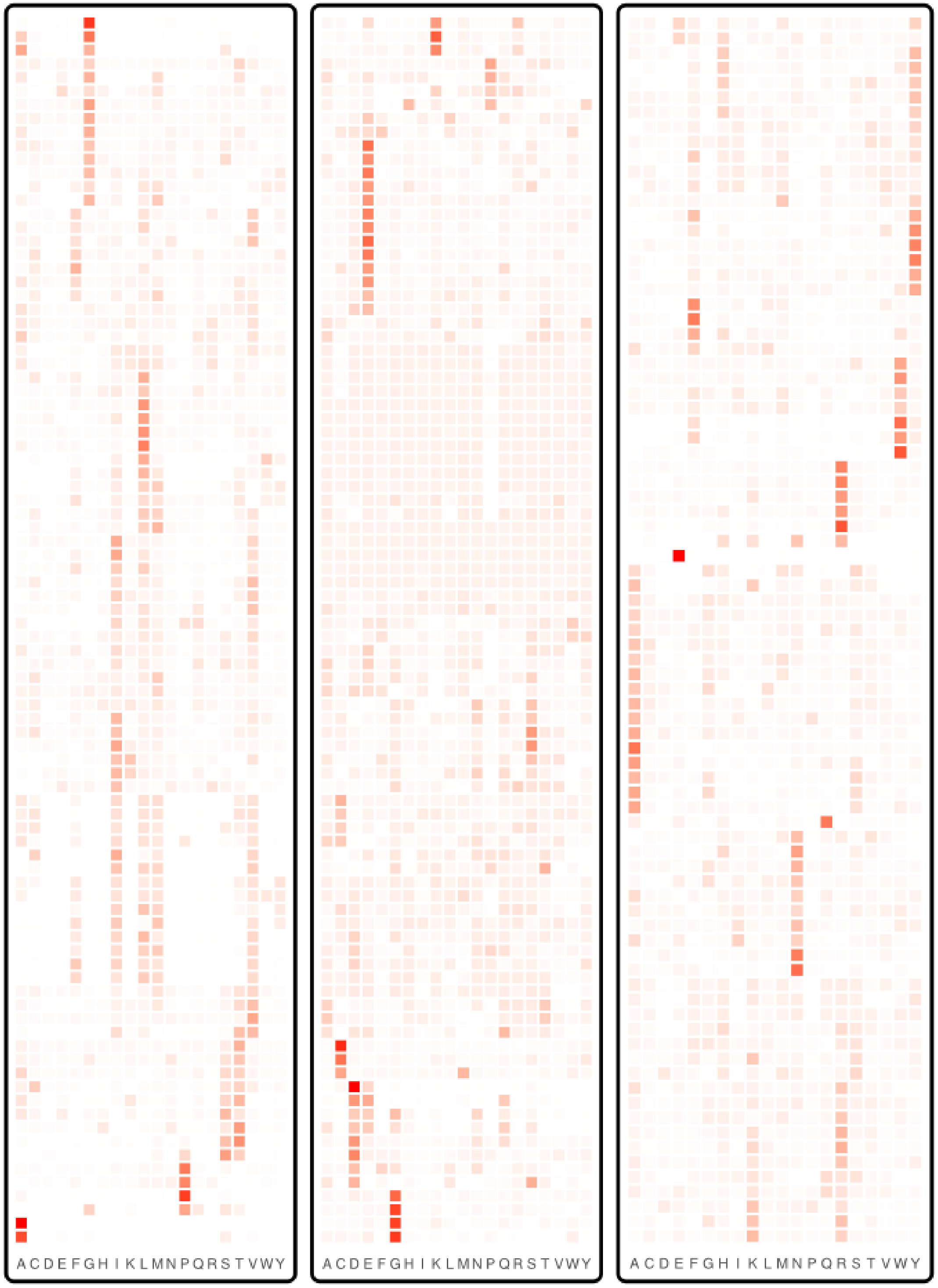


**<u>Figure S2:</u>** Amino acid profiles selected from the Bloom dataset. A. Redundancy in sets of profiles is higher in Bloom dataset than in the 263 retained profiles. Distribution of the distance to the closest profile, measured as the Jensen-Shannon distance. B. The 263 Bloom profiles retained after removing redundancy, ordered by Jensen-Shannon distance. Each row corresponds to a profile and each column to the frequency of an amino acid in this profile. Amino acid frequencies for each profile are available at [https://gitlab.in2p3.fr/pveber/reviewphiltrans](https://gitlab.in2p3.fr/pveber/reviewphiltrans).

### 4/ Choice of NeS used in the simulations

The profiles we used were derived from assays in which the fitness of all 20 amino acids are estimated in competition experiments in the lab, resulting in fitness profiles. Those fitness profiles are not representative of fitnesses that would be obtained in the wild but are highly dependent upon the experimental conditions that were used. In particular, we expect that the scaled selection coefficient NeS would change the fitness (Figure 1). To estimate a realistic range of NeS values to use in our simulations, we computed descriptive statistics on empirical and simulated alignments. We reasoned that NeS values that generate simulated alignments with descriptive statistics similar to those of empirical alignments would be appropriate.

We computed several statistics:

- dS: the number of synonymous substitutions per site summed over the entire tree. Note here that dS is not defined as is typically done in codon models.
- dN: the number of non-synonymous substitutions per site summed over the entire tree. Note here that dN is not defined as is typically done in codon models.
- dN/dS: the ratio of the above two
- Normalized amino acid diversity: the total number of different amino acid states per site, divided by dS

Those statistics were computed by fitting bppml (3) on empirical or simulated codon alignments, and then mapping substitutions using mapnh (3,15). The results were then analyzed using Python and R scripts.

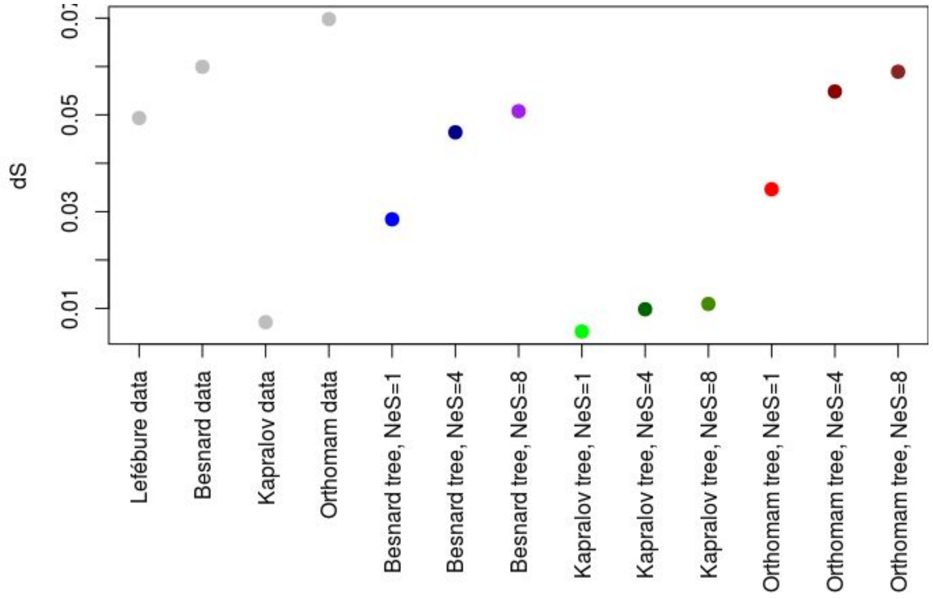


**<u>Figure S3:</u>** dS statistic computed on empirical (4 first columns) and simulated data. The simulated data are based on 3 different tree topologies obtained from the literature (8–10,16), using 3 different NeS values (1, 4, 8), under a homogeneous model and a single amino acid profile per site.

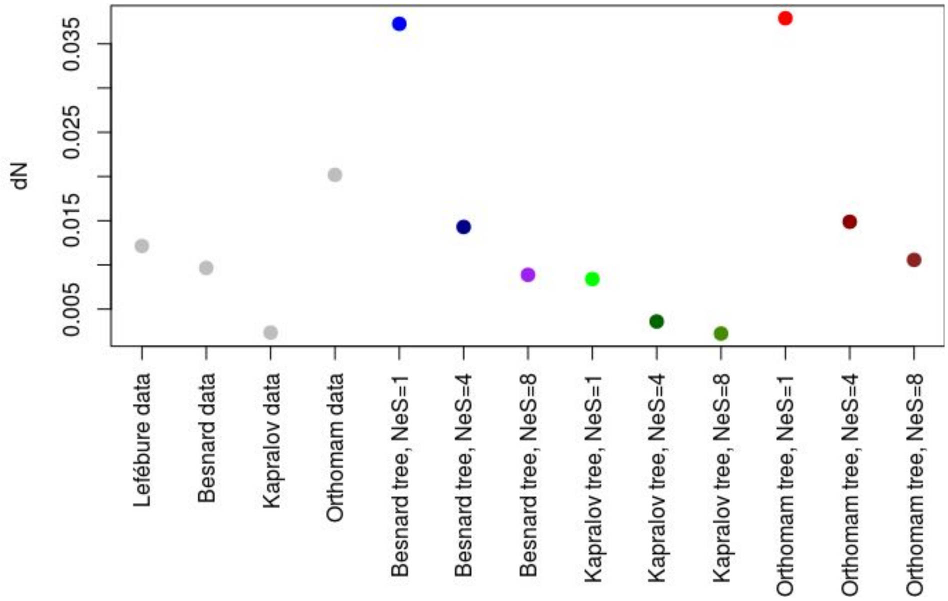


**<u>Figure S4:</u>** dN statistic computed on empirical (4 first columns) and simulated data. Legend is as above.

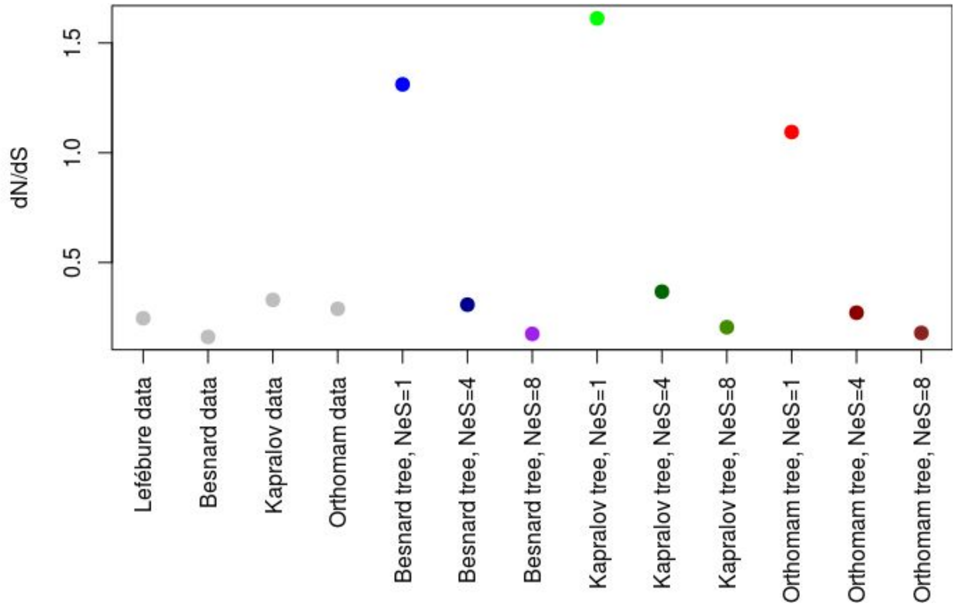


**<u>Figure S5:</u>** dN/dS statistic computed on empirical (4 first columns) and simulated data. Legend is as above.

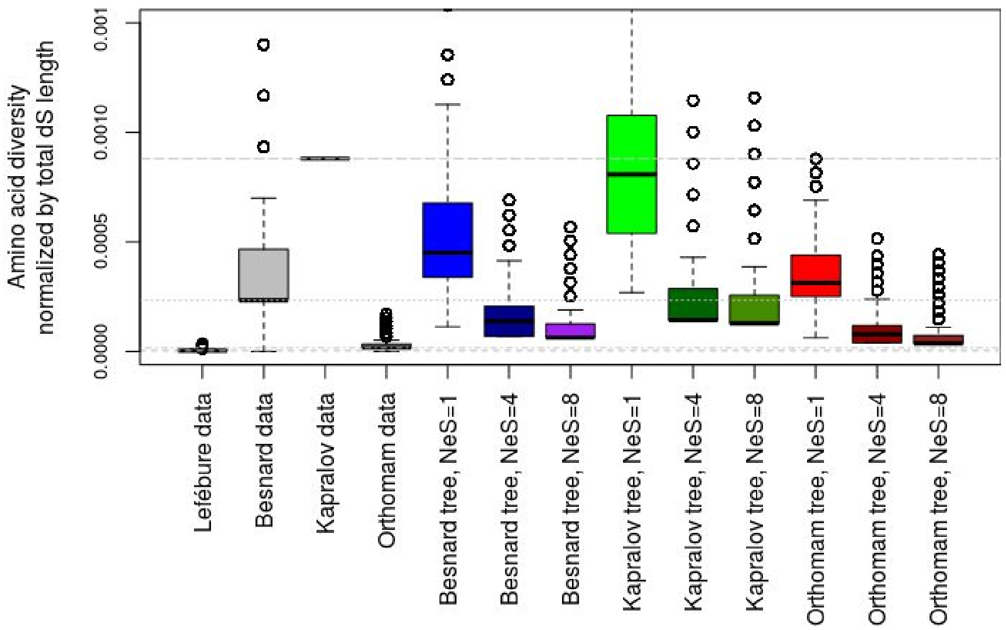


**Figure S6:** Normalized diversity statistic computed on empirical (4 first columns) and simulated data. Legend is as above.

Based on the figures above, we considered that the NeS values 4 and 8 in particular provided data sets that were representative of what is observed in nature.

#### 5/ Standardizing the output of detection methods

In order to compare results across methods, it was necessary to standardize their output. We have decided to use a score between 0 and 1 for every site, where higher means the site is more likely to be convergent.

- For the **Identical** method, the score of a site corresponds to the number of substitutions to the most frequent amino acid found among species with the convergent phenotype. Amino acid frequencies are calculated using bppancestor from (17) using LG08 as substitution model.
- For the **Topological** method, the score of a site is the posterior probability between 2 models using the same parameters except the tree topology, where in the H0 hypothesis, the true topology is used and in the Ha hypothesis a convergent topology where all convergent leaves where group together as defined in (18,19)) and implemented in (20). The likelihood of each model is calculated by bppseqgen from (17) using LG08 as substitution model.
- For **TDG09**, the score of a site correspond to (one minus) the p-value of the likelihood ratio test calculated by the method. We can not use the FDR values given by the method because sites from the H0 and the Ha hypothesis are independently scored which is incompatible with the calculation of the FDR.
- The **diffsel** method, published in (21), uses a codon-based differential-selection model to detect patterns of convergent evolution. A MCMC approach is used to compute

posterior distributions of parameter values, from which statistics can be gathered to establish convergence scores. The original method looks at the probability p(i, aa) over the posterior distribution, for site i and amino acid aa, that the differential selection effect for aa is greater than the mean over all amino acids. If this probability is close to 0 or to 1 (i.e., above 1-c or below c, for some cutoff c<0.5), then it is considered that there is a statistically significant differential effect for this site and amino acid. In order to compare diffsel with other methods, we compute s(i, aa) = 2 * |0.5 - p(i, aa)| to get a score between 0 and 1 such that higher is better. We then compute diffsel_max(i) = $max_{aas}$(s(i, aa)), the maximum score on all amino acids, to get a score per site. Another thing to note is that diffsel is not completely site-independent as it estimates branch lengths across sites. Running it on separate 100% Ha and 100% H0 alignements may thus give skewed results compared to a more realistic case with 2% Ha and 98% H0 in the same alignment. However, early experiments showed (see supplementary section 6), that in practice the difference was small. We thus decided to keep Ha and H0 alignements separate, as alignments with 98% H0 and 2000 Ha sites would have been prohibitively expensive to process (several months per alignment).

- For **PCOC**, the score of a site is the value of the PCOC model given by the method.
- For the **multinomial** method, the score of a site corresponds to 1 - p-value, using the p-value of a Chi2 test to compare the amino acid frequency vector for the extant species with the ancestral phenotype to that obtained for the extant species with the convergent phenotype.
- For **msd**, the score of a site corresponds to 1 - p-value given by the method.

### 6/ Comparaison of the AUC values for the 4 phylogenies

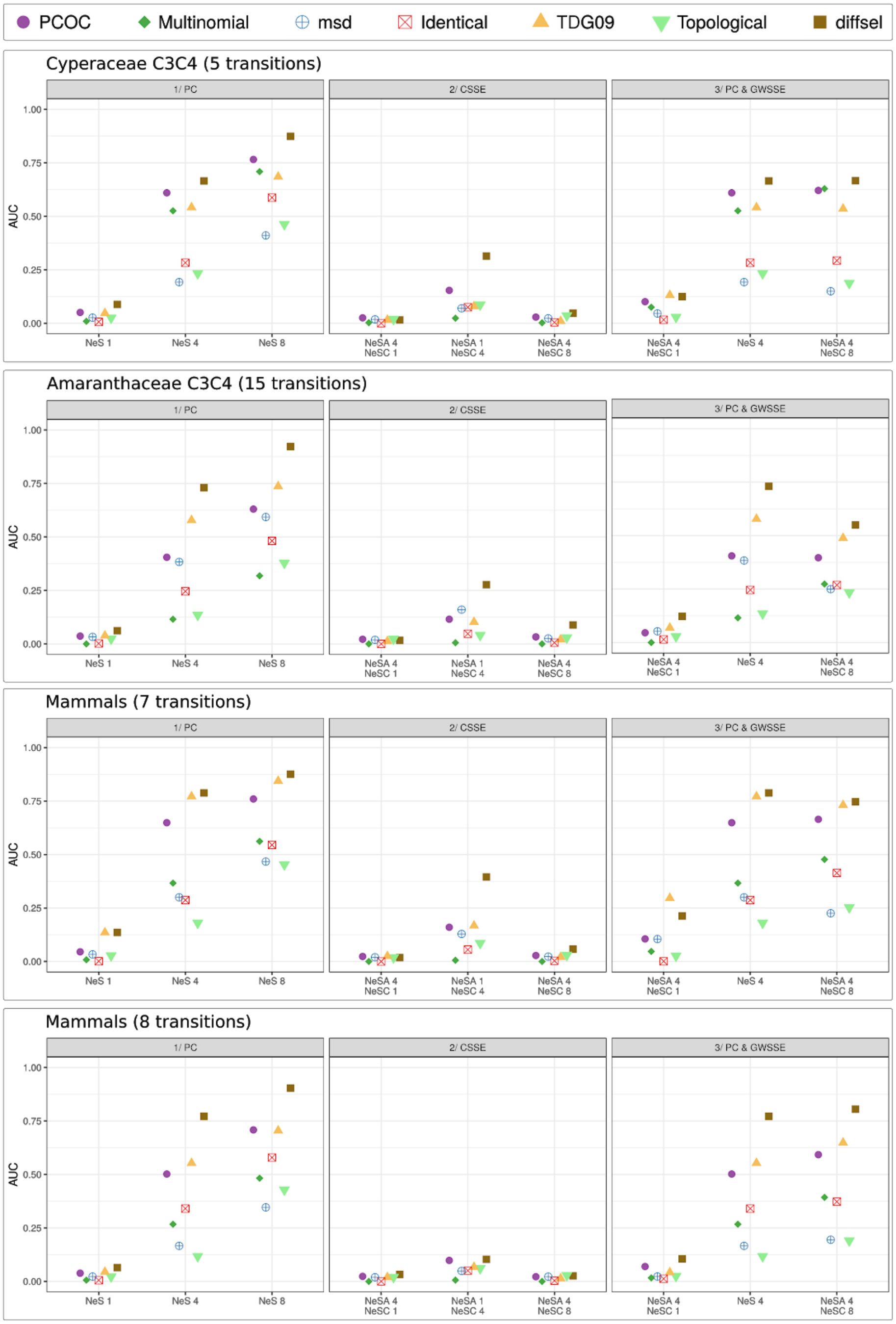


**Figure S7:** Comparison of the AUC values for the 4 phylogenies in the 3 kinds of adaptive convergent cases that have been simulated: 1) a convergent profile change in the selective pressure, 2) a convergent scaling of selection efficacy, and 3) a convergent profile change combined with a selection efficacy scaling. Cyperaceae row corresponds to results presented in Figure 4. The genome-wide selection efficacy ($NeS_A$) remains the same in 1) and is changed to a convergent selection efficacy ($NeS_c$) in Ha 2) and Ha 3).

### 7/ Impact of using separate alignments for diffsel

Diffsel is the only method among the ones we considered that is not site-independent. Indeed, it estimates branch lengths across sites. Running diffsel on separate H0 and Ha alignments may lead to different branch lengths between alignments, making scores difficult to compare across alignments.

In order to estimate the impact of using separate alignments, we have conducted a small experiment outside of our main pipeline where we run diffsel on three different alignments:

- a 900-site H0 alignment simulated on the Cyperaceae C3C4 tree with NeS=4;
- a 100-site Ha alignment simulated with profile change and the same tree and NeS;
- the 1000-site concatenation of the two first alignments.

The concatenated alignment allows diffsel to estimate branch lengths across a 90/10 mix of Ha and H0 sites, without knowledge of which is which. This corresponds to a real alignment with a few convergent sites and a lot of non-convergent ones. We have considered in the paper that, in real data, the H0/Ha ratio would probably be closer to 98/2. Running diffsel on an alignment with such a H0/Ha ratio would, however, require a very large alignment in order to keep a reasonable number of Ha sites.

Figure S8 presents the results of these runs in the form of recall/precision plots corrected to mimic a 98/2 H0/Ha ratio, like in the paper. Table S1 gives figures for AUC and best recall at 90% precision. We see that the concatenated runs have slightly worse results in terms of AUC, but still stay ahead of PCOC, the second best method in the same conditions. Recall at 90% precision stays the same. Overall, assuming a similar behaviour across NeS hypotheses, using diffsel on concatenates would not change any of the qualitative results presented in the paper, but would have been extremely compute-intensive. Indeed, we would have needed to simulate 2000 Ha sites, which means 2000*98/2=98000 H0 sites so as to run Diffsel in conditions similar to the other methods. Such a run would take several months with the current (single-threaded) diffsel implementation.

**Table S1:** AUC and best recall at 90% precision for diffsel on 900/100 H0/Ha concat and non-concat simulations. AUC for pcoc in similar conditions (from Figure 3) is given for comparison.

| | **Separate** | **Concat** |
|---|---|---|
| **AUC diffsel_max** | 0.707 | 0.621 |
| **R90 diffsel_max** | 0.51 | 0.51 |
| **AUC pcoc** | 0.609 | |

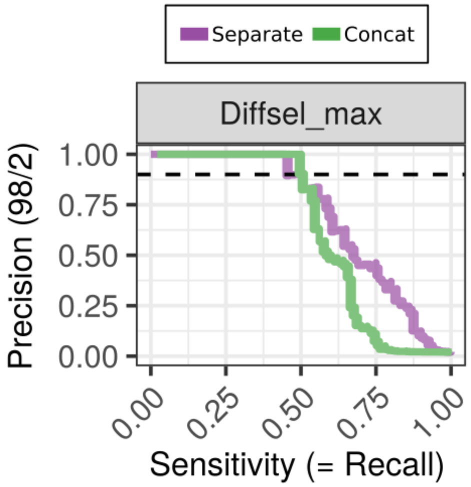


**Figure S8:** Precision/recall plots for diffsel on separate alignments (purple) or concatenated alignments (green).

The results presented in Figure S8 have been computed by running diffsel and scripts manually outside of the pipeline. Intermediate results and reproduction steps are available at ftp://pbil.univ-lyon1.fr/pub/lanore/diffsel_concat_experiment.tar.gz (300Mo) during the review process (and would be hosted on Dryad after acceptance).

### 8/ How to reproduce our results

Apart from the observed computing times for detection methods, results in this paper should be exactly reproducible.

**Computing resources:** keep in mind that the whole pipeline can take a very long time to run depending of the number of trees, the number of hypotheses and the number of simulated sites. Experiments presented in the paper took five days to run on a 24-core machine for 2000 sites, 4 trees and 12 hypotheses. Suitable computing resources should be used to reproduce our run (at least 24 cores for five days or 48 for two days and a half). Running all the experiments on a 4-core laptop would take approximately a month.

**Installed software:** a working version of docker is necessary to run our pipeline. Installation instructions for docker are beyond the scope of this reproduction guide. Be aware that docker requires root privileges and is thus unlikely to be available on traditional clusters. Virtual machines are an easy way to circumvent this problem.

Apart from docker, opam (an OCaml package manager) and a few OCaml packages are necessary to run the pipeline. The following commands should install all of that on a Debian-based linux distribution:

```
sudo apt install m4 opam zlib1g-dev
opam init
eval `opam config env`
opam switch 4.06.1
eval `opam config env`
opam pin add biocaml https://github.com/biocaml/biocaml.git
```

```
opam pin add bistro https://github.com/pveber/bistro.git#first-class-file-dumps -n
opam install ppx_csv_conv bistro ocamlify biocaml
```

All other software is packaged in docker images and thus requires no installation. The use of docker images guarantees that the various programs in the pipeline (scripts, detection methods, python and R packages…) are exactly identical to the ones we used in our experiments.

**Pipeline version:** the pipeline itself is available through the repository at https://gitlab.in2p3.fr/pveber/reviewphiltrans. The version that we used for the paper corresponds to tag "v1.01" (commit d3e25255c8315a29a81a9cc8a691005568dee57e).

**Build and run the pipeline:**

```
git clone https://gitlab.in2p3.fr/pveber/reviewphiltrans
cd reviewphiltrans
git checkout v1.01
make
make install
make analyses
```

**Basic troubleshooting:** an error at compilation (make) probably indicates that opam or some packages are not properly installed: try re-running the opam commands given above. An error at runtime mentioning docker probably indicates that docker is improperly installed or configured.

**Pipeline output:** if the pipeline successfully runs, a folder will be created called `reviewphiltrans/example/outdir_analyses`. This folder contains all intermediate (e.g., simulated data, method outputs, .tsv files used for plots…) and final (plots) results. Results are organized in sub-folders with explicit names. The plots used in the paper are called `<tree-name>.recall_precision_ok.pdf`.

**Paper intermediate results:** all the intermediate results of the runs used in the paper have been conserved to aid reproducibility. They are temporarily available at ftp://pbil.univ-lyon1.fr/pub/lanore/outdir_full.tar.gz (16Go) during the review process (and would be hosted on Dryad after acceptance). A lighter version (500Mo) of the intermediate results without the diffsel traces (which take 800Mo per run) is available at ftp://pbil.univ-lyon1.fr/pub/lanore/outdir_without_chains.tar.gz.
